# Breaking the “unbreakable” ZIKA virus envelope

**DOI:** 10.1101/345884

**Authors:** Chinmai Pindi, Venkat R Chirasani, Mohd. Homaidur Rahman, Mohd. Ahsan, Prasanna D Revanasiddappa, Sanjib Senapati

## Abstract

The rapid spread of zika virus (ZIKV) and its association with microcephaly and Guillain-Barre syndrome have raised major concerns worldwide. Studies have shown that ZIKV can survive in harsh conditions, *e.g.* high fever; in sharp contrast to the dengue virus (DENV) of same Flaviviridae family. In spite of recent cryo-EM structures that showed similar architecture of the ZIKV and DENV envelopes, little is known of what makes ZIKV envelope so robust and unique. Here, we present a detailed analysis of the constituent raft-raft and protein-protein interactions on ZIKV and DENV envelopes to undermine their differential stability at near-atomic to atomic level using coarse-grained (CG) and all-atom (AA) molecular dynamics simulations. Our results from CG simulations show that, at high temperatures, ZIKV envelope retains its structural integrity, while DENV2 disintegrate through the formation of holes at 5- and 3-fold vertices. Protein structural network from AA simulations shows a stronger inter-raft communications in ZIKV through multiple electrostatic and H-bond interactions. Particularly, the intricate network of interlocking DE-loop and FG-loop among five DIII domains in ZIKA vertices was exceedingly robust that makes this envelope stable, even at high temperatures. Our results are validated by alanine mutations to the CD-loop residues Gln350 and Thr351 that showed no effect on ZIKA stability, in close accordance with a recent mutagenesis study. These detailed information, which were difficult to extract experimentally, broadened our understanding of the flaviviruses and can accelerate the structure-based drug designing processes aiming ZIKA and DENV therapeutics.

**IMPORTANCE:** The rapid spread of zika virus (ZIKV) and its association with severe birth defects have raised worldwide concern. Recent studies have shown that ZIKV can survive in harsh conditions, *e.g.* high fever, unlike dengue (DENV) and other flaviviruses. Here, we unravel the molecular basis of ZIKV unprecedented stability over DENV at high temperatures, mimicking fever. Our study, based on coarse-grained and all-atom molecular dynamics simulations, could not only explore the mechanism of DENV envelope breaking at high temperatures, but also captured atomic-level contacts and interactions at raft-raft and protein-protein interfaces that keep ZIKV envelope intact in similar conditions. The obtained results are validated by *in-silico* and reported *in-vitro* mutagenesis studies by showing the presence of specific H-bonding and electrostatic interactions among ZIKV E protein residues. At the end, our study was successful to define potential ZIKV epitopes and provide residue-level insights for designing specific ZIKV antibodies and small molecule inhibitors.

## INTRODUCTION

The recent spread of zika virus (ZIKV) and its association with microcephaly and Guillain-Barre syndrome have raised major concerns worldwide. Since its recent report in 2015 from Brazil, the virus has affected more than a million individuals across the world.(1) Consequently in March 2016, WHO declared an international health emergency over zika virus outbreak that caused serious birth defects. The zika virus is transmitted to human primarily by mosquitoes, but reports of transmission through other means have also been documented recently.(2, 3) The virus belongs to the family of flaviviruses that also includes dengue, west nile, japanese encephalitis, and yellow fever viruses.(4)

Recent cryo-EM studies have revealed that the morphology of zika virus (ZIKV) envelope (5, 6) is very similar to that of the other flaviviruses of known structure, such as the dengue virus (DENV).(7) The viral envelope comprises of 180 copies of E and M proteins that arrange in an icosahedral geometry. While the surface-exposed E proteins are crucial in receptor binding and fusion, the hidden M proteins primarily anchor into the lipid membrane covering the viral RNA. The E protein consists of 504 amino acids, with the N-terminal 403 residues forming the ectodomain that folds mainly in β-sheets. Two of these E proteins assemble in the form of head-to-tail homodimers, with three such homodimers lying parallel to each other to form a flat-boat like surface known as raft (Fig. 1). There are 30 such rafts which constitute the whole viral envelope. Also, each E protein is comprised of three domains: DI, DII, and DIII with domain DI bridging between domains DII and DIII (Fig. 1C). The intersection of five rafts at the periphery of five DIII domains constitutes a 5-fold vertex, while the intersection of three rafts at the periphery of three DI domain constitutes a 3-fold vertex on the virus envelope surface(5) (Figs. 1A, 1B). There are twelve 5-fold and twenty such 3-fold vertices on the ZIKV/DENV envelope.

**Fig 1:**
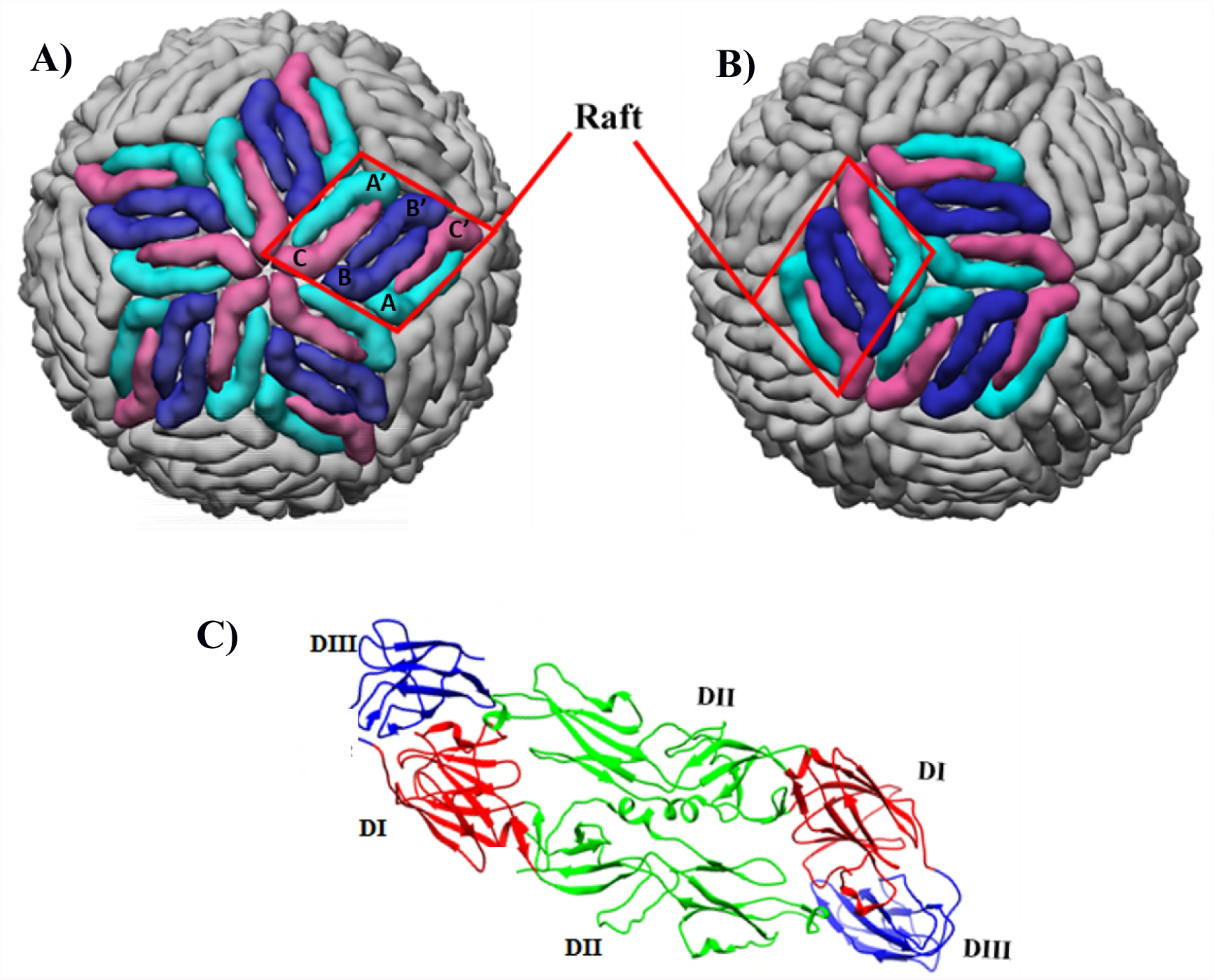
Architecture of Icosahedral viral envelope. Viral envelope surface highlighting (A) a 5-fold vertex at the intersection of five neighboring rafts (labeled) and (B) a 3-fold vertex at the intersection of three neighboring rafts. Conventional nomenclature of the six constituent E proteins in a raft is used with molecule A/A’ shown in cyan, molecule B/B’ in blue, and molecule C/C’ in pink. **(C)** E protein homodimer showing the constituent domains.

Although the cryo-EM studies have revealed similar envelope structures of zika and other flaviviruses,(6) the more recent reports based on various biological assays have suggested that only ZIKV can sustain the harsh conditions of saliva, urine, semen etc.(8). Even more interestingly, unlike other flaviviruses, ZIKV showed very little sensitivity to high temperatures. For example, when exposed to a temperature of 37°C, Fibriansah et al. found significant expansion in the E protein shell of dengue virus with a hole at its 3-fold vertices.(9) Later, Lim et al. found serotype-specific expansion in this virus, with the DENV serotype 2 (DENV2) exhibiting notable instability in its structure at 37°C(10) On the contrary, Kostyuchenko et al. (6) shown that the incubated ZIKA samples at 37°C, and even at 40°C (temperature mimicking high fever), displayed smooth surfaced particles in the electron microscopy maps, while the dengue particles were broken and shrivelled at both these temperatures. In consistence with this finding, plaque assay shows little change in infectivity of ZIKV, whereas the infectivity of DENV was greatly affected with increasing temperature.(6) The structural basis of this unusual stability of ZIKV over DENV is unknown. In this study, we attempt to understand the underlying mechanism of the differential stability of ZIKV and DENV2 at varying temperatures.

Even though the cryo-EM studies have provided important information about the structures of different flavivirus envelopes, the atomistic details pertaining to their differential stability is yet to be known. Here, we employ coarse-grained (CG) and all-atom molecular dynamics (MD) simulations to explore both the long and short time and length scale phenomena occurring on the virus envelopes, since the envelope constitutes the first level of protection to the viral RNA and reported to be majorly responsible for viral stability. The state-of-the-art computational techniques employed here can not only explore the mechanism of virus envelope breaking, but also can capture atomic-level contacts and interactions at protein-protein interfaces that are difficult to trace experimentally.

The first successful attempt of utilizing MD simulation techniques to explore the structure and dynamics of virus particles was by Schulten and coworkers. In consistence with AFM and other biophysical studies, these authors showed pronounced instability of the satellite tobacco mosaic virus (STMV) capsid in absence of its RNA core.(11) In yet another pioneering MD study, these authors have shown that brome mosaic virus capsid, poliovirus capsid, the bacteriophage øX174 procapsid, and the reovirus core could be stable without their respective genetic material.(12) These studies have provided plethora of atomic-level details that have improved our understanding of the viruses immensely. Since then different groups have modelled and simulated various enveloped and non-enveloped viruses, such as HIV, influenza A, Rous sarcoma virus, HBV etc.(13–15) Very recently, Reddy et al. have performed coarse-grained MD simulations of DENV2 envelope including the lipid bilayer to understand the viral protein-lipid interactions.(16) Marzinek et al. presented a refined model of DENV2 envelope anchored with a more realistic membrane composition that shows greater morphological resemblance with the cryoEM constructs.(17) Thus, the robustness of MD simulation methods in expanding the horizon of our understanding of viruses is well documented. However, ZIKA virus that has become a serious threat to the next generation human population, is yet to be investigated at this detailed level. Here, we report such an in-depth study of ZIKV for the first time to unravel the unique network of interactions present on its envelope surface that help the virus sustain harsh conditions, such as high temperatures in comparison to DENV2.

Our results show that at temperatures mimicking high fever, the envelope of ZIKV was intact while the DENV2 envelope disintegrated through the raft-raft interfaces, triggered by the formation of holes at 5- and 3-fold vertices. Stronger energetics at raft-raft interfaces showed the presence of multiple polar and H-bonding interactions in ZIKV compared to DENV2. Finally, we identified a set of key ZIKV residues that play crucial role in its stability, which upon mutation destabilized the virus envelope. Our study provides useful insights to specifically target the ZIKA virus envelope for designing novel antiviral therapeutics.

## RESULTS AND DISCUSSION

To start with, we performed CG simulations of hollow and water-filled envelopes of both ZIKV and DENV2 at 29°C and 40°C (Table S1). Subsequently, more sophisticated all-atom MD simulations were executed. Results show an unprecedentedly large number of polar and H-bonding interactions across the ZIKV raft-raft interfaces, which were absent in DENV2. Mutations of these interfacial residues destabilized the ZIKV envelope significantly. The detailed results are presented in the following sections.

### Stable ZIKV envelope at 40°C

The multi-microsecond long CG simulations reproduced the cryo-EM results by showing greater thermal stability of the zika virus over the dengue. At ambient temperature of 29°C, both the virus envelopes were stable. However as the temperature was increased to 40°C, the ZIKV envelope was intact while DENV2 broke through the 5-fold vertices. To quantify the observed changes, we first computed the root mean square deviations (RMSD) of each of the 30 rafts that constitute the virus envelope. RMSD of a raft was calculated by monitoring the displacements of the constituting six E protein backbones with respect to their conformations in the cryo-EM map. As the results in Figs. S1A-B indicate, both the virus envelopes were stable at 29°C. All the constituting 30 rafts exhibited a narrow range of RMS deviations from the respective EM structures. When the virus particles were filled with water, the RMSD values reduced further due to favourable interactions between the solvent and the protein residues of the rafts (Figs. S1C-D). Notably, while all rafts were stabilized equally in the water-filled envelope of ZIKA, a small set of DENV rafts continued to exhibit similar RMSD values as that in the hollow envelope at ~ 0.8 nm, while rest others better stabilized at 0.4 nm. The relatively large RMSD values of the virus rafts are not unexpected considering the fact that each raft is composed of six asymmetric units which are not covalently bonded to each other, and each asymmetric unit has 1737 residues in ZIKV and 1701 residues in DENV.

As the temperature was increased to 40°C, the ZIKA envelope remains equally compact as it was at 29°C. This was evident from the togetherness of the RMSD plots of all the 30 rafts. Both the hollow and water-filled simulated systems represent well-stabilized ZIKA envelope, where the RMSD plots reach to a plateau within the first 500 ns of simulation time (Figs. S1E, 2A). On the contrary, a set of DENV rafts exhibited a sharp increase in RMSD values within the first 50 ns of simulation time (Figs. S1F, 2B), suggesting loosely-packed DENV envelope that could be disrupting at high temperature.

To confirm the above finding of compact *versus* loosely-packed envelope of ZIKA *versus* DENV, we also examined the time variation of raft-raft angles. The raft-raft angle was calculated by defining two vectors – Â and Ĉ that respectively connect the center of mass (COM) of the envelope to the COM of the most stable raft (*i.e.,* the raft with the least RMSD, termed as “base raft”) and the COM of the envelope to the COM of any other raft (Fig. 2C). The angle between the two vectors, q was then computed as follows:

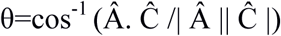

**Fig 2:**
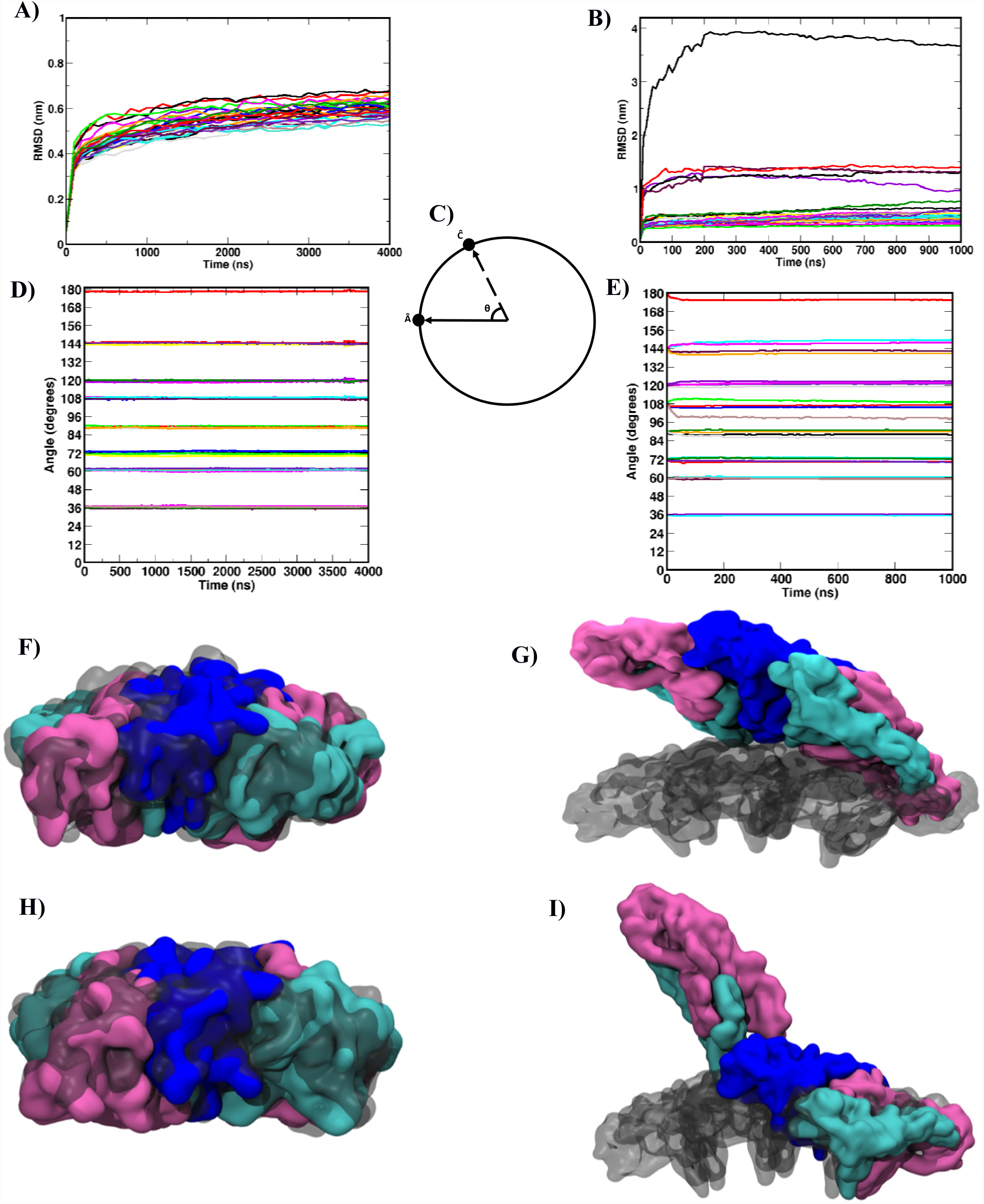
Stable ZIKV *versus* unstable DENV envelope at 40°C. Time evolution of RMSD of each of the 30 rafts that constitute the virus envelope in **(A)** ZIKV^40^_water_ and (**B**) DENV^40^_water_ systems from CG simulations. **(C**) Schematic representation of the measured angle (θ) between two rafts (see text for details). Time evolution of the angle between “base raft” and each other rafts in **(D)** ZIKV^40^_water_ **(E)** DENV^40^_water_ systems. Two representative DENV rafts **(G, I)** that undergone significant changes from their starting conformations are shown. Color scheme of the six constituent E proteins is similar to Fig. 1A. Starting conformations of the rafts are shown in grey. Corresponding ZIKV rafts **(F, H)** show minimal changes as evident from their perfect superposition on the starting conformations.

The time evolution of the inter-raft angles at 29°C for both ZIKA and DENV2 envelopes show that seven raft assemblies, with each assembly consisting of four rafts remain stable throughout the simulations at their respective native angle of 36°, 60°, 72°, 90°, 108°, 120°, and 144°, relative to the base raft (Figs. S2A-D). Further, the single raft located right opposite to the base raft in the envelope remains stable at an angle of 180° across the simulation time. Interestingly, a similar stability was displayed by ZIKA envelope at the elevated temperature of 40°C, as shown in Figs. 2D and S2E for the water-filled and hollow systems, respectively. On the other hand, the inter-raft angle distributions for both hollow and water-filled DENV2 systems at this temperature exhibited clear disruption of the envelope with the raft assemblies segregating apart, suggesting the asymmetric movements of the rafts and possible breaking of the envelope in DENV (Figs. 2E, S2F).

A closer look into the RMSD and raft-raft angle distributions of DENV envelope at 40°C revealed that the five rafts, which underwent the major displacements during simulation constituted a 5-fold vertex in the native envelope (*i.e.* in the cryo-EM structure). Figs. 2F-I show the variations of two of these rafts that exhibited the most significant displacements. While the first DENV raft displaced from the native angle of 144° to 149° (Fig. 2G), the switched from 108° latter to 115° (Fig. 2I). Such an out of phase displacement of two neighboring rafts could implicate a plausible breaking of the DENV envelope. We will elaborate on this in next sections. The superposed 3D representations of the rafts in Figs. 2G and 2I also suggest that while the first DENV raft shows only a change in orientation, the latter shows a breaking at the dimer-dimer interface in the raft. Astonishingly, in spite of such an array of large changes in DENV, at this high temperature all ZIKA rafts remained extremely stable and exhibited minimal variations from the cryo-EM structure (Figs. 2F, 2H).

### Broken DENV envelope at 40°C

It was evident from the RMSD and inter-raft angle calculations that the arrangements of E proteins on ZIKV envelope was very tight, while the assembly in DENV envelope was weak. The weak inter-raft interactions could not sustain the high temperature and thus the DENV envelope disarrayed. Fig. 3 shows the mechanism of breaking of DENV envelope at 40°C. It’s noteworthy that while the DENV particle broke within 50 ns of simulation time, ZIKA remained completely intact at this high temperature even after a much longer simulation time of 4µs. Fig. 3 is produced from the angle of a 5-fold vertex that initiated the breaking of DENV envelope, as traced from our MD simulation trajectory (see Movies S1, S2). As Fig. 3 and Movies S1-S2 show, the ZIKA envelope maintained its compactness (Figs. 3A-B), while DENV enlarged through the protrusion of one of its 5-fold vertices (Figs. 3C-F). To start with, this DENV vertex was as closed as ZIKA (Fig. 3A-C). However, as the simulation progressed, the DENV raft-raft interactions at the 5-fold vertex weakened and a hole is produced (Fig. 3D). At around 15ns, the interactions became weaker and the hole enlarged (Fig. 3E). At 30 ns, the neighbouring 3-fold vertices also protruded out and the DENV envelope lost its icosahedral symmetry (Fig. 3F). Notably, apart from the increased raft-raft separations, some of the rafts exhibited dimer-dimer breaking as discussed in Fig. 2I and shown in Fig. 3F. For the rest of the 1µs simulation, the structure remained similarly or more broken confirming the instability of DENV virus at high temperatures.

**Fig 3:**
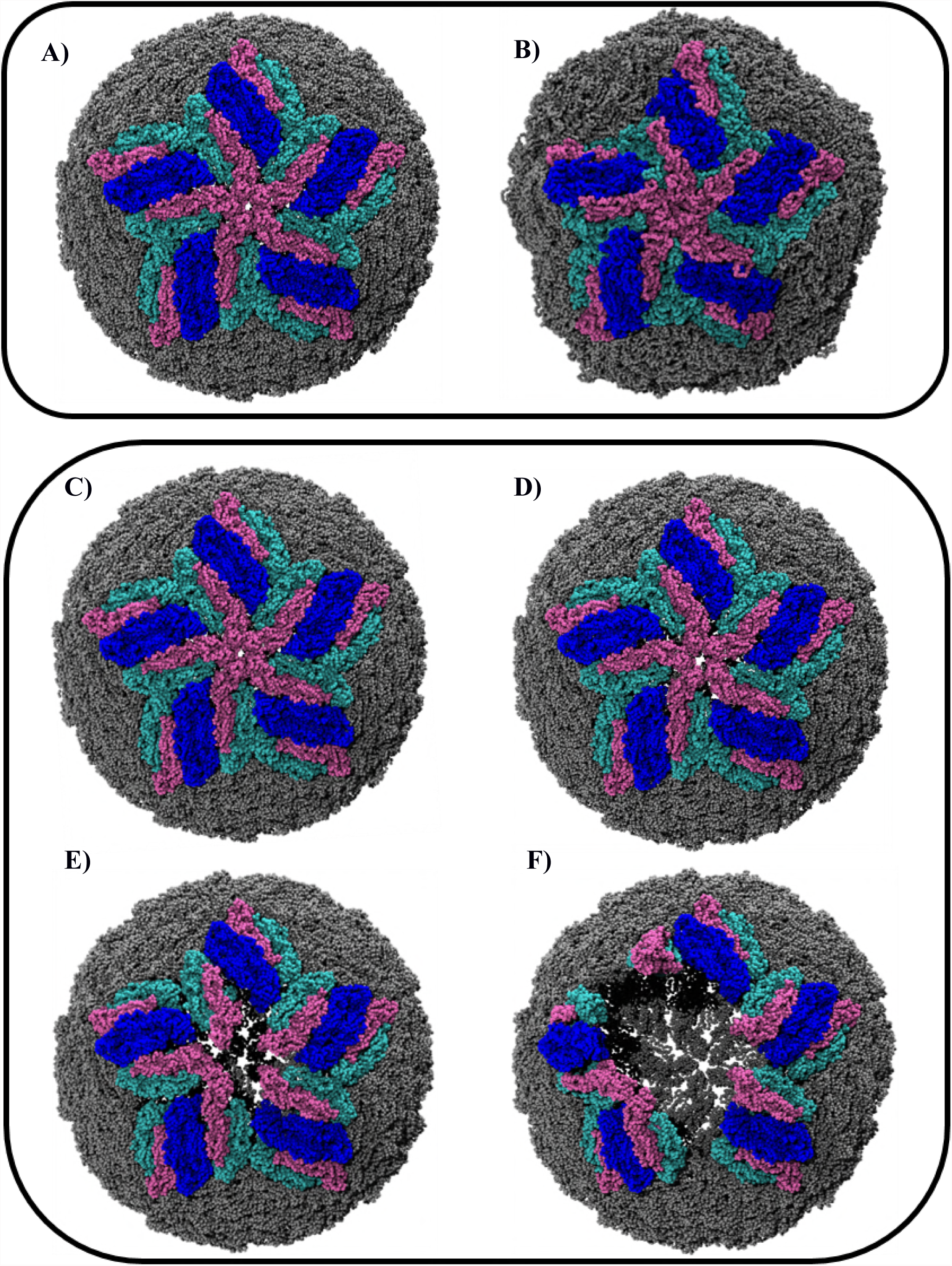
Compact and smooth surfaced ZIKA *versus* loose and rough surfaced DENV envelope. Structure of ZIKV40_water_ system at **(A)** 0ns and **(B)** 4µs of CG simulation time. Structure of DENV^40^_water_ system at **(C)** 0ns, **(D)** 4ns, **(E)** 15ns and **(F)** 50ns of simulation time. Color scheme is similar to Fig. 1A.

Our mechanism matches very well with that of Fibriansah et al., who explored structural changes in dengue virus through cryo-electron microscopy at 37°C.(9) In agreement with different class of DENV2 particles defined by these authors, time evolution of the envelope breaking in our MD simulation captured all class I – III DENV particles (Fig. 4). As the time evolution of DENV envelope radius shows (Fig. 4B), our starting structure mimicked the class I particle with a radius of about 240 Å. As the simulation progressed, the envelope gradually enlarged to 250 Å corresponding to class II and further to 260 – 280 Å, resembling the class III DENV particle of Fibriansah et al. In agreement with the cryo-EM maps, our results also show the loss in icosahedral symmetry of the DENV envelope through the protrusions of its 5- and 3-fold vertices as discussed in Fig. 3. This enlargement consequently led to the loss in structural compactness as manifested in the increased radius of gyration of the viral envelope (Fig. 4B). As we will see shortly, this was associated with the change in surface morphology from smooth to bumpy, primarily due to the protrusion of domain III of C molecules of the E protein at the 5-fold vertices. Such a close resemblance with the experimental data validates our simulation protocol.

**Fig 4:**
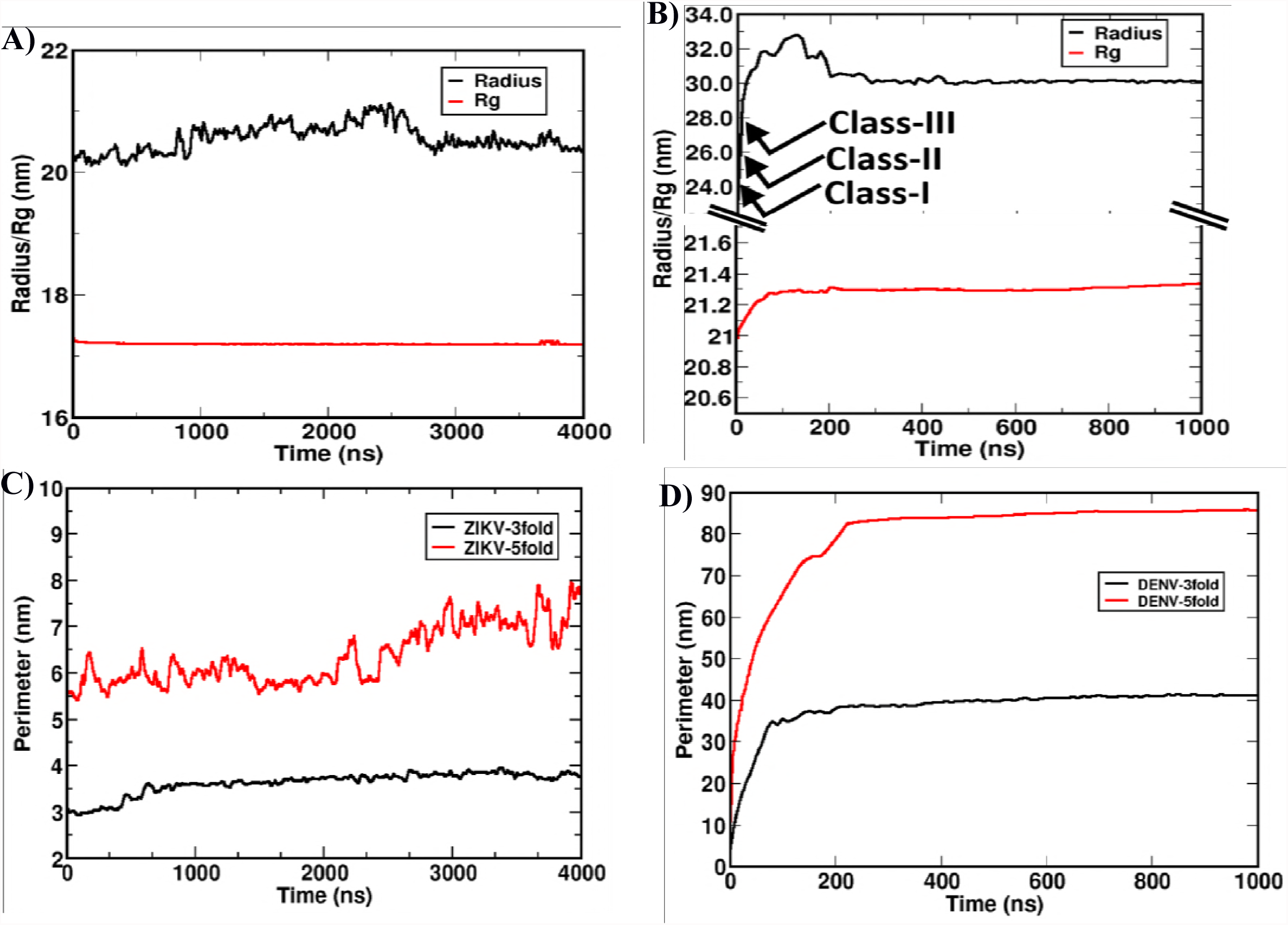
Breaking of DENV envelope nucleates at 5-fold vertices. Time evolution of the radius (black) and radius of gyration (red) of the viral envelope in **(A)** ZIKV^40^_water_ **(B)** DENV^40^_water_ systems from CG simulations. In (B), class I - III correspond to broken DENV envelope of increasing radius. Time evolution of the perimeters at the most deformed 3-fold (black) and 5-fold (red) vertices in **(C)** ZIKV^40^_water_ and **(D)** DENV^40^_water_ systems.

Along the similar lines of RMSD and inter-raft angles, the size and compactness of ZIKA envelope remained almost unchanged during the whole simulation period. As shown in Fig. 4A, both the radius and R_g_ of water-filled ZIKA envelope at 40°C remained very close to the native values. However, the stability of this envelope was best described by quantifying the perimeters of its 5- and 3-fold vertices. As shown in Fig. 4C, both the vertices maintained the average perimeter values that are very close to the cryo-EM structure. The plotted perimeters were the running averages over all the 12 and 20 number of 5- and 3-fold vertices present in the envelope. On the contrary, both the perimeters in DENV envelope increased manifold from the native structure values. Fig. 4D shows the time-dependent opening of the specific 5-fold vertex that initiated the breaking of the envelope. While comparing the rate of opening of this 5-fold vertex with the neighbouring 3-fold vertex that exhibited the most significant opening among all 3-fold vertices, it was observed that the kinetics of breaking at the 5-fold vertex was faster than the 3-fold. The slope at 10-50 ns time span was 0.54 *versus* 0.35 for the respective 5- and 3-fold vertices. This led us to conclude that the instability of dengue particles at high temperatures is initiated by the breaking of 5-fold vertices on its envelope structure, which subsequently triggers the disintegration of 3-fold and other inter- and intra-raft regions.

To strengthen the above findings, we performed new sets of CG simulations of water-filled ZIKV and DENV envelope both at 29°C and 40°C (Table S1). In these simulations, a different set of initial velocities of the constituent atoms were chosen randomly from a Maxwell-Boltzmann distribution. Results from these replica simulations also revealed a temperature-insensitive ZIKV *versus* temperature-sensitive DENV envelope structure (Fig. S3). More importantly, these simulations also displayed a similar mechanism of DENV breaking through the 5-fold vertices that subsequently extends to 3-fold vertices and other regions through the weakening of inter-and intra-raft contacts.

From the above discussion it is evident that ZIKV envelope retains its structural integrity, while the DENV2 envelope breaks at high temperature. Although the CG simulations provided sufficient clue for this differential behaviour of the two homologous virus envelopes, the atomic-level details could not be obtained due to the low-resolution trajectories from CG simulations. Hence in the second half of the work, we employed united atom (UA) simulations to compare the ZIKV and DENV envelopes at finer details (Table S1). In the UA simulations, except for the non-polar hydrogens, each other atoms of the virus envelopes were described explicitly. The non-polar hydrogens were embedded with the heavy atoms to which they were bonded. Similar to the CG simulations, the UA simulations were also initiated from the cryo-EM structures of ZIKV and DENV.

### Compact and smooth surfaced ZIKA *versus* loose and rough surfaced DENV envelope

We carried out 20 ns long UA simulation for each of the ZIKA and DENV envelope at 40°C. It is worth mentioning here that simulating these large systems of more than 12 million atoms requires extensive computational time, and the runtime for 1 ns simulation in our available resource of 256 cores Intel Xeon E5-2650 processors varied from 48 - 52 hrs. Hence, simulating these systems for more than 20 ns was difficult. Nevertheless, as shown below, this time length was sufficient to extract new insights, since the results converged to EM and CG simulation findings. These atomic-level information, which were difficult to extract experimentally, could broaden our understanding of the flavivirus family and also can accelerate the structure-based drug designing processes aiming ZIKA and DENV therapeutics.

To start with, we examined the stability of the 5-fold vertices in each system as it was the pivot of DENV envelope breaking. Similar to the cryo-EM and CG simulation results, ZIKV displayed a compact structure while the DENV envelope exhibited significant instability. This has been shown by comparing the time-averaged perimeters of the 5-fold vertices of the viral envelopes. As shown in Fig. 5, all the twelve 5-fold vertices in ZIKA envelope remain very stable and their perimeters deviate very little from the cryo-EM structure. In fact, majority of the 5-fold vertices in ZIKV have shown a decrease in perimeter with one of the vertex contracting as low as 51 Å compared to the EM value of 59 Å, demonstrating the compactness of the envelope. The average of all 12 ZIKA perimeters was 55.11 Å. On the contrary, the vertices in DENV were greatly distorted and the perimeters increased significantly from the reference EM structure value. Most of the vertices enlarged with one of them expanding as big as 69 Å compared to the EM value of 55 Å, signifying drastic loosening up of envelope structure. The average value of all 12 DENV perimeters was 64.84 Å. Thus, the widened 5-fold vertices in DENV envelope confirms its swollen state at high temperature from atomistic simulation. Moreover, regular *versus* deformed pentagonal shapes of the 5-fold vertices (compare Fig. 5A with Fig. 5B) signify smooth *versus* rough ZIKA and DENV particles, as observed experimentally. It is also worth mentioning here that, similar to the CG simulations data, UA simulations also suggested that 5-fold vertices nucleate the DENV envelope breaking, as exemplified by its faster increase in perimeters than the 3-fold vertex perimeters (Fig. S4).

**Fig 5:**
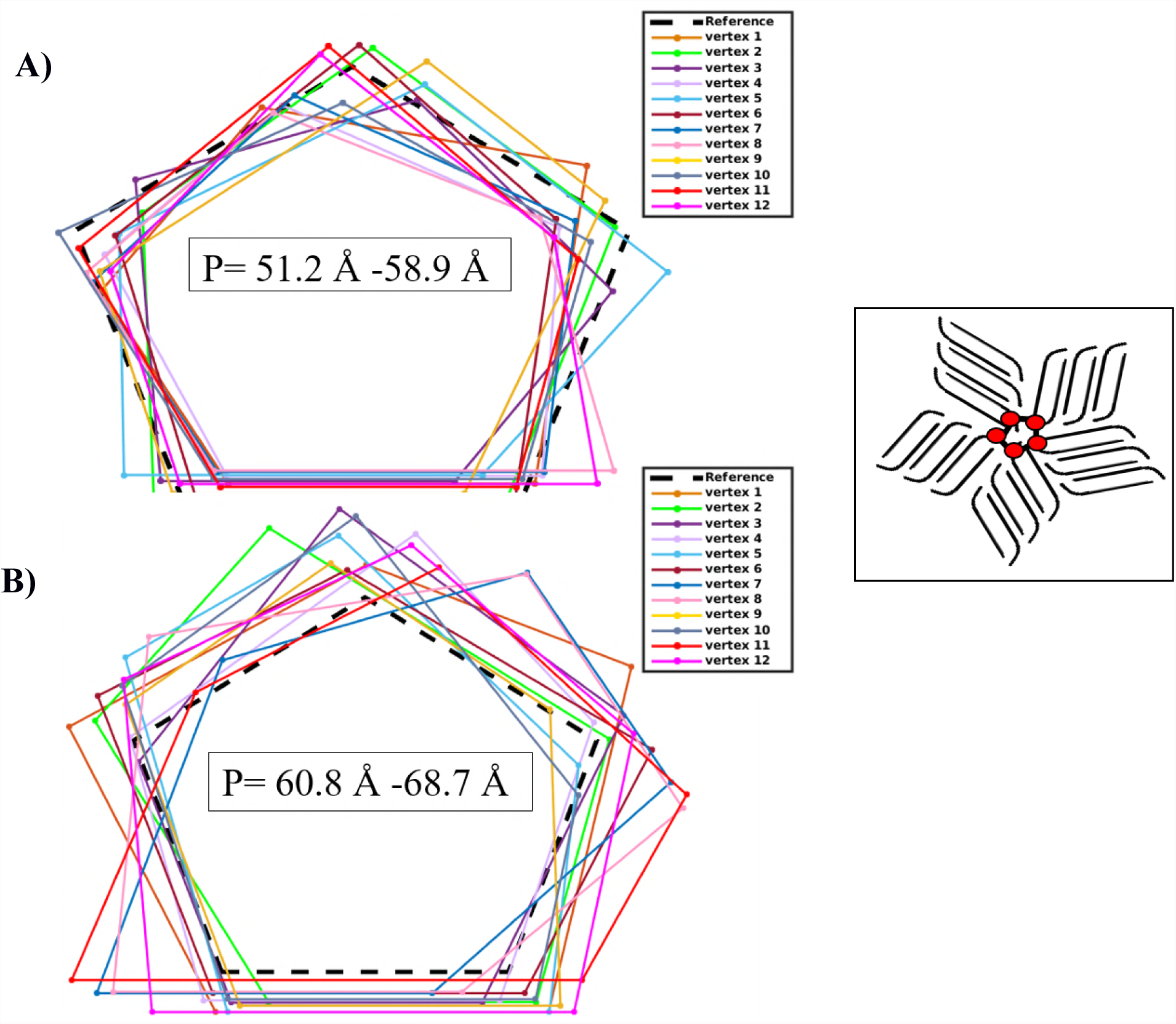
ZIKV 5-fold vertices remain compact at high temperature. Time-averaged perimeters of all the twelve 5-fold vertices in **(A)** ZIKV^40^_water_ and **(B)** DENV^40^_water_ systems from UA simulations. The calculated perimeters from EM structures are included in black dotted lines. The span of all 12 perimeters in each envelope is shown. Inset shows the schematic diagram of a 5-fold vertex that attains regular pentagonal shape in EM structure.

The vertex perimeters in Fig. 5 were calculated by summing up five center of mass (COM) distances of the FG-loops (7) in ZIKV and DENV. These loops belong to DIII domain of E protein C molecule and five such loops from five individual C molecules encompass a 5-fold vertex in the viral envelope structure. These loops are almost equidistant from each other in the EM structure and observe a regular pentagonal shape at the vertex, as shown in the schematic diagram in Fig. 5 inset. The plotted distances in Fig. 5A-B are the time-average over 20 ns of the simulation trajectory. It is also to be noted from Fig. 5 that vertex 11 in DENV (shown in red) underwent the highest deformation. This observation was confirmed from the largest RMS displacement experienced by this vertex from the EM structure, out of all 12 vertices constituting the envelope (Fig. S5). In the subsequent discussion, therefore, we offer a special emphasis on the changes experienced by this representative vertex and its constituent rafts.

### Stronger inter- and intra-raft contacts in ZIKV

Fig. 6A depicts the two-dimensional representation of the most deformed 5-fold vertex in ZIKA at 40°C from the UA simulation. As evident, the vertex experienced negligible changes with the five constituting rafts maintaining strong inter-raft contacts. To quantify this, we calculated the number of contacts at each of the five raft-raft interfaces and the results are shown in Fig. 6B. The results suggest that the number of contacts in all five interfaces in ZIKV remains almost fixed, while in DENV the contacts reduced significantly. There was a total decrease of 8327 number of contacts (or 20.8% decrease) in DENV, compared to a 10.1% total increase in ZIKA. Notably, while three of the interfaces in DENV exhibited a reduction of 40%, 23% and 31% contacts compared to their respective values in the native form, the other two interfaces showed about ~10% increased contacts. This increased contacts can easily be understood by looking at Fig. 3 and Movie S2, which showed that breaking at one interface brings the two other adjoining rafts somewhat closer. The loss of contacts at DENV inter-raft interfaces primarily stems from the weaker interactions of the FG-loops of DIII domain at 5-fold vertices. At inter raft region, the interaction of CD-loop with bc-loop (DII) and eE_0_-loop (DI) of adjacent rafts were also diminished.(7) Apart from this, weakening of interactions between hi-loop (DII) and l_0_A-loop (this loop is at the linker region of DI and DIII) from the neighbouring rafts were also noted in DENV. On the other hand, these interactions were intact in ZIKV that exhibited rather an increased inter-raft contacts, in consistent with its decreased 5-fold perimeter shown in Fig. 5a.

**Fig 6:**
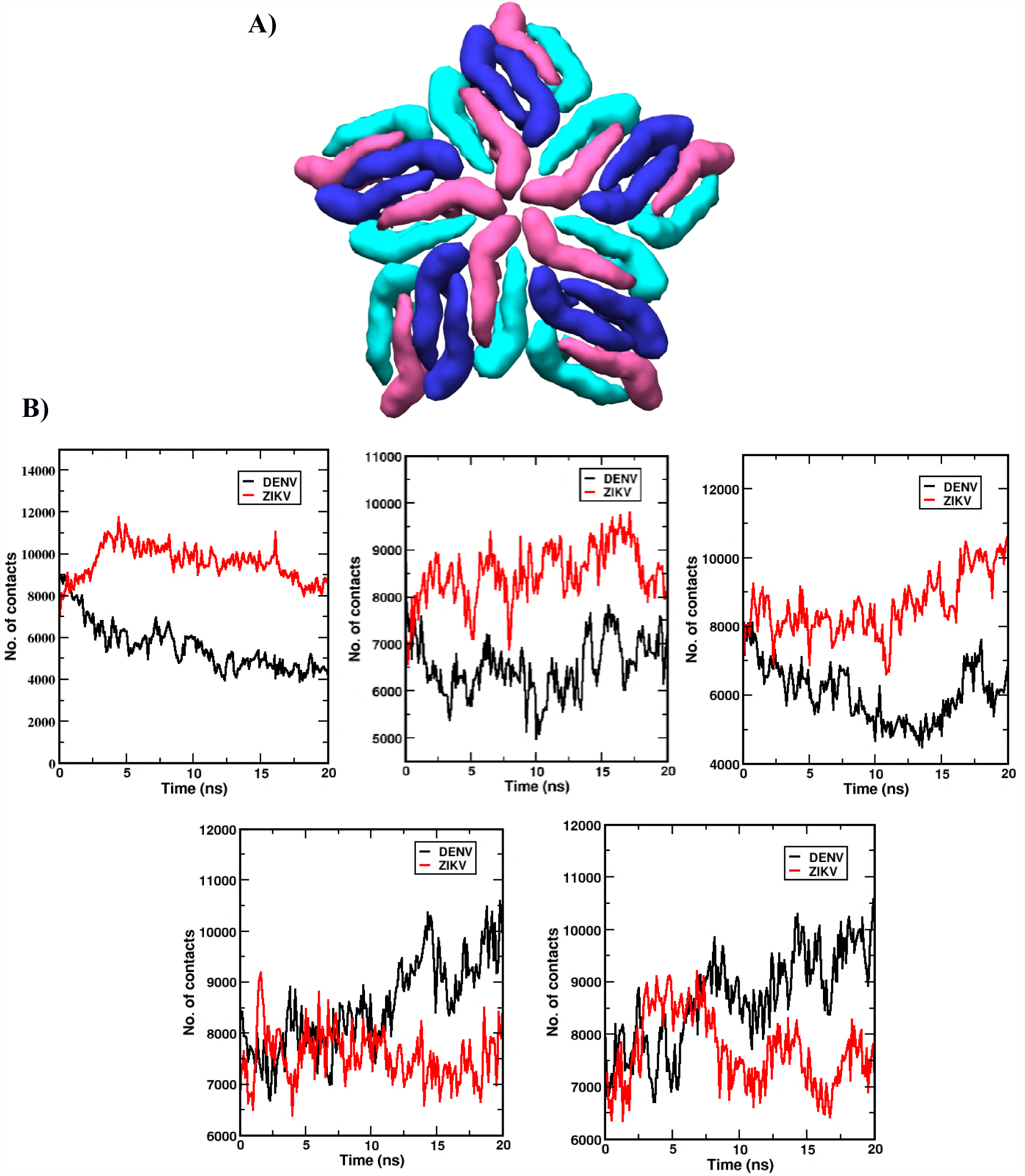
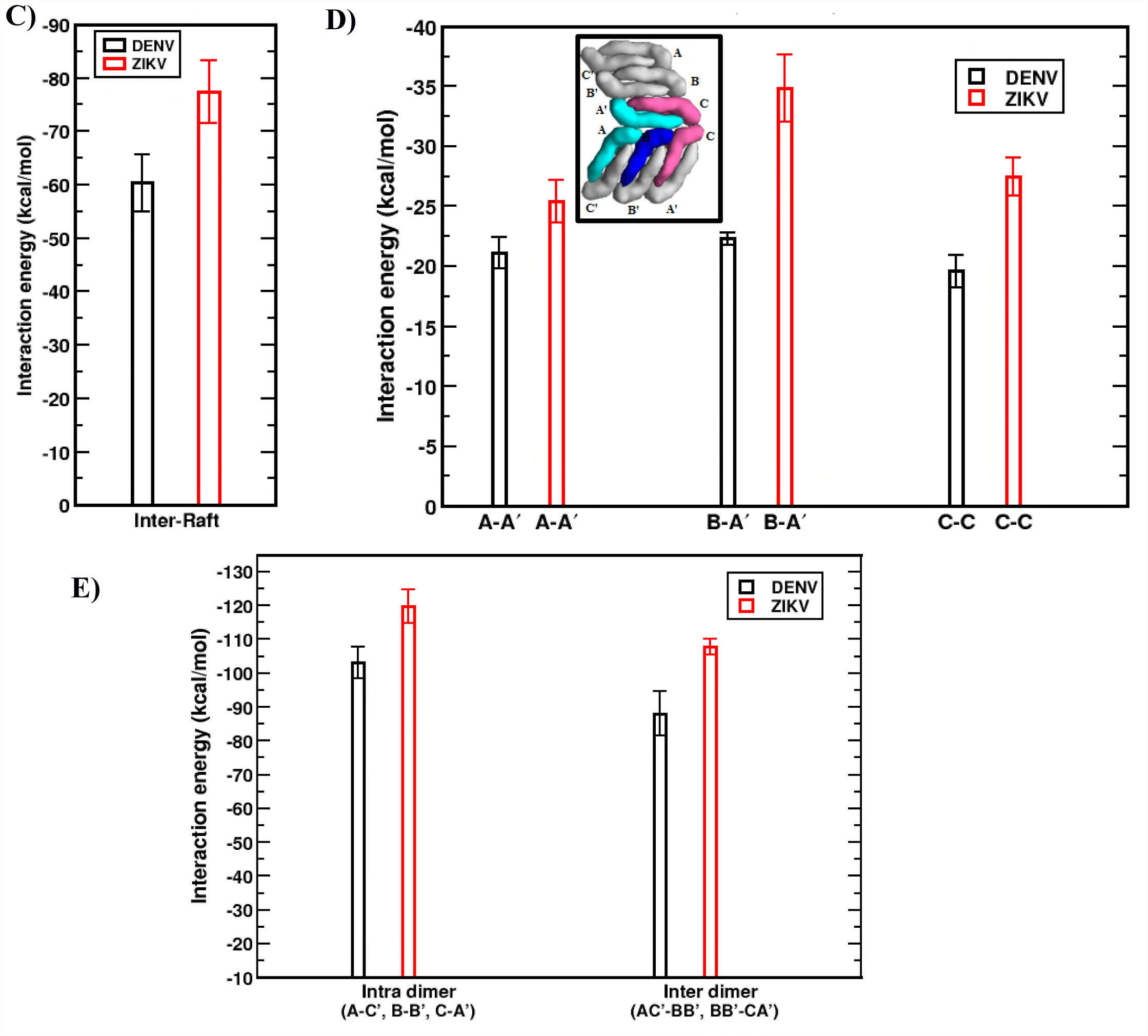
Stronger inter- and intra-raft contacts in ZIKV than DENV. **(A)** The 2D map of a representative 5-fold vertex in ZIKV at 40°C from UA simulation. Color scheme is similar to Fig. 1A. **(B**) Comparison of the number of contacts at five inter-raft interfaces present in the 5-fold vertex shown in (A). **(C)** Average inter-raft interaction energy in ZIKV^40^_water_ (red) and DENV^40^_water_ (black). **(D)** Average interaction energy between two E protein molecules at the ZIKV (red) and DENV (black) inter-raft interfaces. Inset shows the distributions and specific interactions between the E protein molecules from two neighboring rafts. **(E)** Average inter- and intra-dimer interaction energy in ZIKV (red) and DENV (black). See Fig. 6D inset for E protein notations.

To make a direct comparison of the raft-raft interactions in ZIKA and DENV, we then calculated the average raft-raft interaction energy over the five inter-raft interfaces present at the representative 5-fold vertex shown in Fig. 6A. The energy values were computed over the first 10 ns simulation data, since the DENV envelope started exhibiting instability beyond this point. As Fig. 6C shows, the average raft-raft interaction energy was significantly larger in ZIKA (−78.36 kcal/mol) compared to DENV (−60.37 kcal/mol). To find the respective contributions of the constituent E protein molecules, we subsequently split the total raft-raft interaction energy into protein-protein interaction energy. As Fig. 6D inset shows, the interactions at a raft-raft interface is primarily contributed by the association of constituting protein molecule A from one raft with protein molecule A’ of the adjacent raft (A-A’), association of protein molecule B of one raft with protein molecule A’ of the adjacent raft (B-A’), and association between protein molecule C of the first raft with protein molecule C of the second raft (C–C). Figure 6D shows the comparison of these protein-protein interactions at ZIKV and DENV inter-raft interfaces. It is evident that protein-protein interactions in ZIKV is much greater than DENV with a energy difference of ~3 kcal/mol at A-A’ interface, ~11 kcal/mol at B-A’ interface, and ~7 kcal/mol at C-C interface. In spite of similar interactions at DENV A-A’ and C-C interfaces, the greater susceptibility of DENV envelope breaking through the C-C interface (5-fold vertex) might stem from its less buried surface area (1300Å^2^) compared to the A-A’ interface (1500 Å^2^). The presence of more buried surface area results in a more disordered solvent environment that increases the solvent entropy and hence total free energy at the A-A’ interface in DENV.(18, 19)

We also looked into the intra-raft association by calculating the inter-dimer AC’-BB’, BB’-CA’ and intra-dimer A-C’, B-B’, C-A’ interaction energies (see Fig. 6D inset for the protein molecule nomenclature). Results in Fig. 6E suggest that intra-raft interactions - both inter-dimer and intra-dimer – are not too different in ZIKA and DENV. However, the noteworthy features that emerge from Figs. 6D and 6E are - (i) inter-raft interfaces are always weaker than the intra-raft interfaces, (ii) intra-dimer association is the strongest among all the protein-protein interactions present, in both ZIKA and DENV. These results are consistent with above CG simulation results showing the breaking of DENV particle through the raft-raft interfaces and the experimental report showing that E proteins exist as dimers in solution.(9) Overall, our results suggest that ZIKV has stronger intra- and inter-raft interactions that make this virus envelope stronger than the DENV.

### Electrostatics and H-bonding interactions impart greater ZIKV inter-raft stability

From the above results it is evident that DENV envelope is more susceptible to break at high temperature through its inter-raft interfaces. To find the constituent E protein residues that were responsible for such differential inter-raft stability of ZIKA *versus* DENV, we looked into the residue-level energy contribution at the inter-raft interfaces of both the envelopes. As can be noted from Fig. 6D inset, the stability of A-A’ interface is primarily contributed by the interactions of domain DIII residues of A molecule with the domain DI residues of A’ molecule of the second raft (see Fig. 1C for domain names). Similarly, at B-A’ interface, it is the interaction of DIII domain residues of B molecule with the DII domain residues of A’ molecule of the second raft that were responsible for the stability. The C-C interface which is inferred to play a key role in the stability of the envelope involves the interactions of DIII domain residues of C molecules from the adjacent rafts.

Figures 7A-C show the residue-level energy contributions at the three inter-raft regions - A-A’, B-A’, and C-C in both ZIKA and DENV with the domain-wise sequence alignments of two viral E proteins depicted in the insets. Results are shown for the residue-pairs that contributed ≥ −1 kcal/mol to the total energy of the respective inter-raft interfaces. As Fig. 7A shows, at the inter-raft region A-A’, nearly equal number of residue pairs contribute to the total energy in ZIKV and DENV. However, while majority of these residue pairs were involved in electrostatic and H-bonding interactions in ZIKA, they involve in hydrophobic or weakly electrostatic interactions in DENV. For example, the residue pairs Thr353_r1_-Glu133_r2_, Lys395_r1_-Asn134_r2_, Gln344_r1_-Asn172_r2_ from two neighbouring rafts, r1 and r2 in ZIKA showed > −3 kcal/mol energy contribution, in contrast to the very little contribution of the corresponding residue pairs in DENV. Though the residues Glu133_r2_ and Asn134_r2_ were conserved in domain I of both viral proteins, domain III showed substitution from Thr353_r1_ and Lys395_r1_ in ZIKV to His346_r1_ and Gln386_r1_ in DENV. While Thr353_r1_ could involve in strong sidechain-sidechain H-bonding interactions with Glu133_r2_, and Lys395_r1_ could interact electrostatically with Asn134_r2_ in ZIKA; DENV substitutions fail to sustain such favourable interactions. Similarly, stable H-bond was observed between the side chain of Gln344_r1_ and backbone amide of Asn172_r2_ in ZIKV. H-bond lifetime analysis have shown that both the above H-bonds persisted up to 70% of simulation time (Fig. S6A). On the contrary, no H-bond existed more than 20% of simulation time in DENV. These results also match favourably well with the ZIKA and DENV cryo-EM structure data.(5, 7) For instance, the observed H-bond between backbone amide of Leu352 and sidechain of Glu133 in ZIKA cryo-EM structure was persistent over 50% of the simulation time (indicated by * in Figs. S6A, 7A). On the contrary, the reported H-bonds between Ser298 and Glu172 and between Arg345 and Glu133 in the DENV cryo-EM structure,(5, 7) existed only for 12% and <10% of the simulation time (indicated by # in Figs. S6A, 7A), suggesting significantly weaker interactions in DENV, which further weakens at higher temperature. It is worth mentioning here that some of the aforementioned H-bonds in the cryo-EM structures were observed upon modelling the missing atoms/residues.

**Fig 7:**
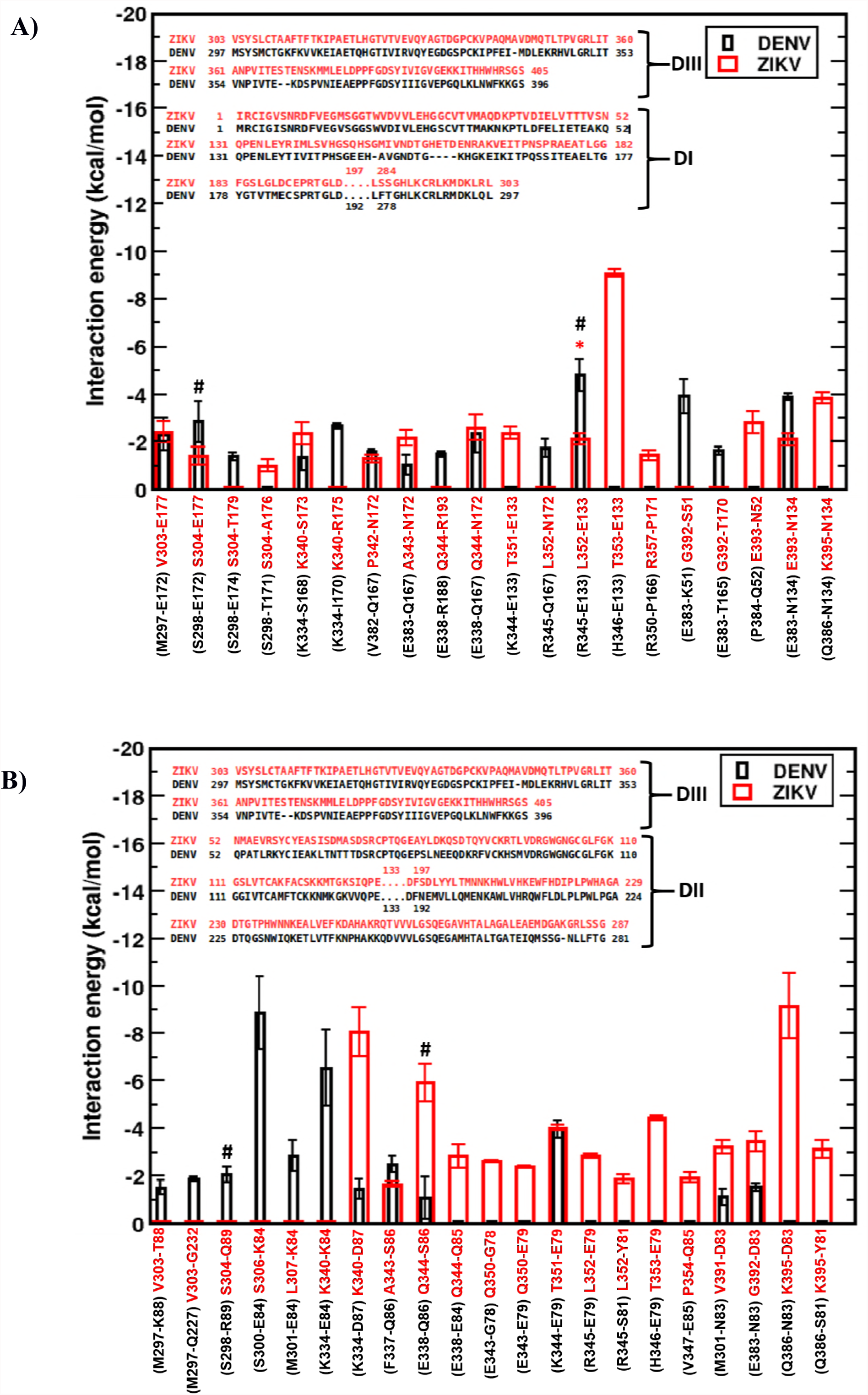
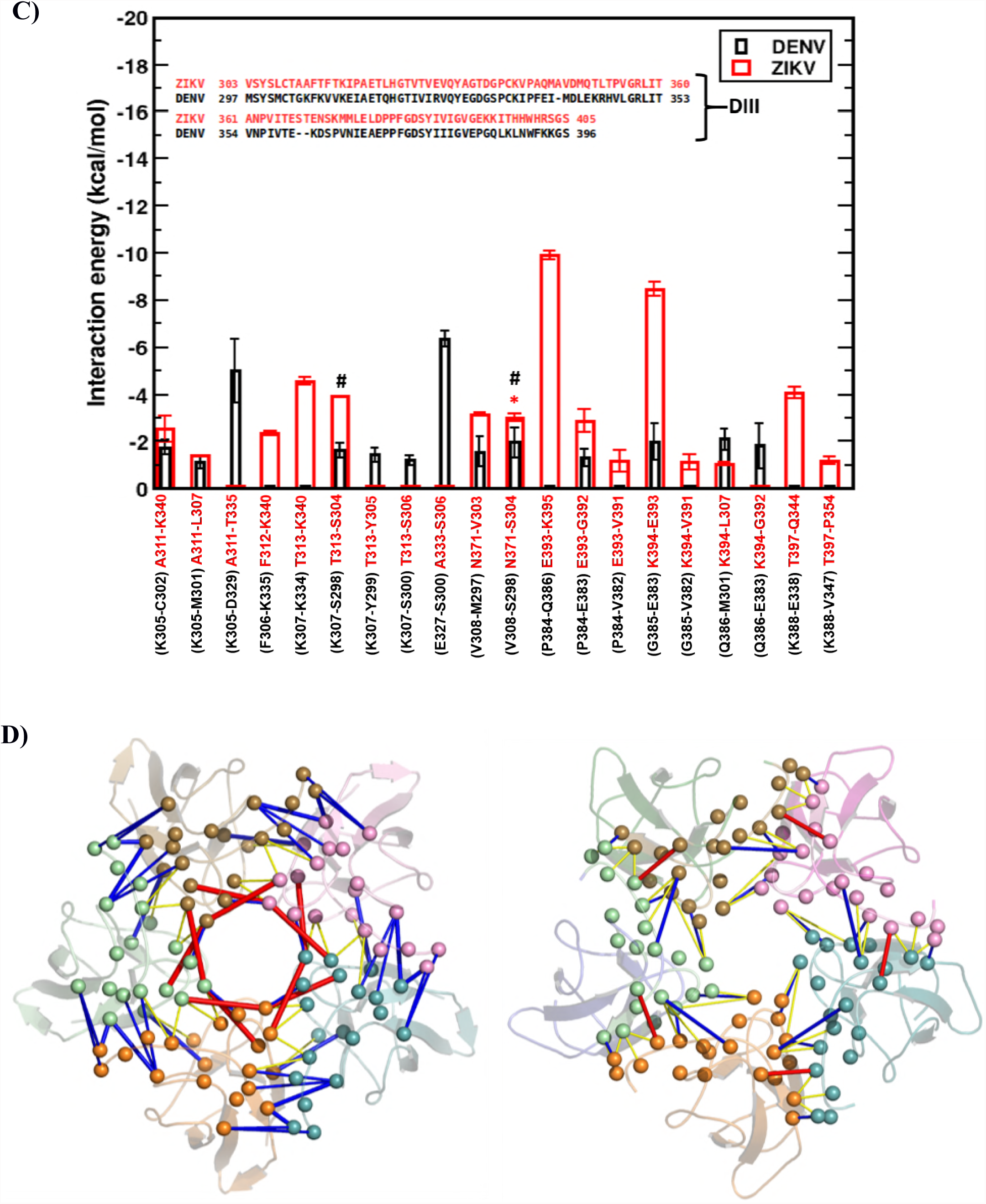
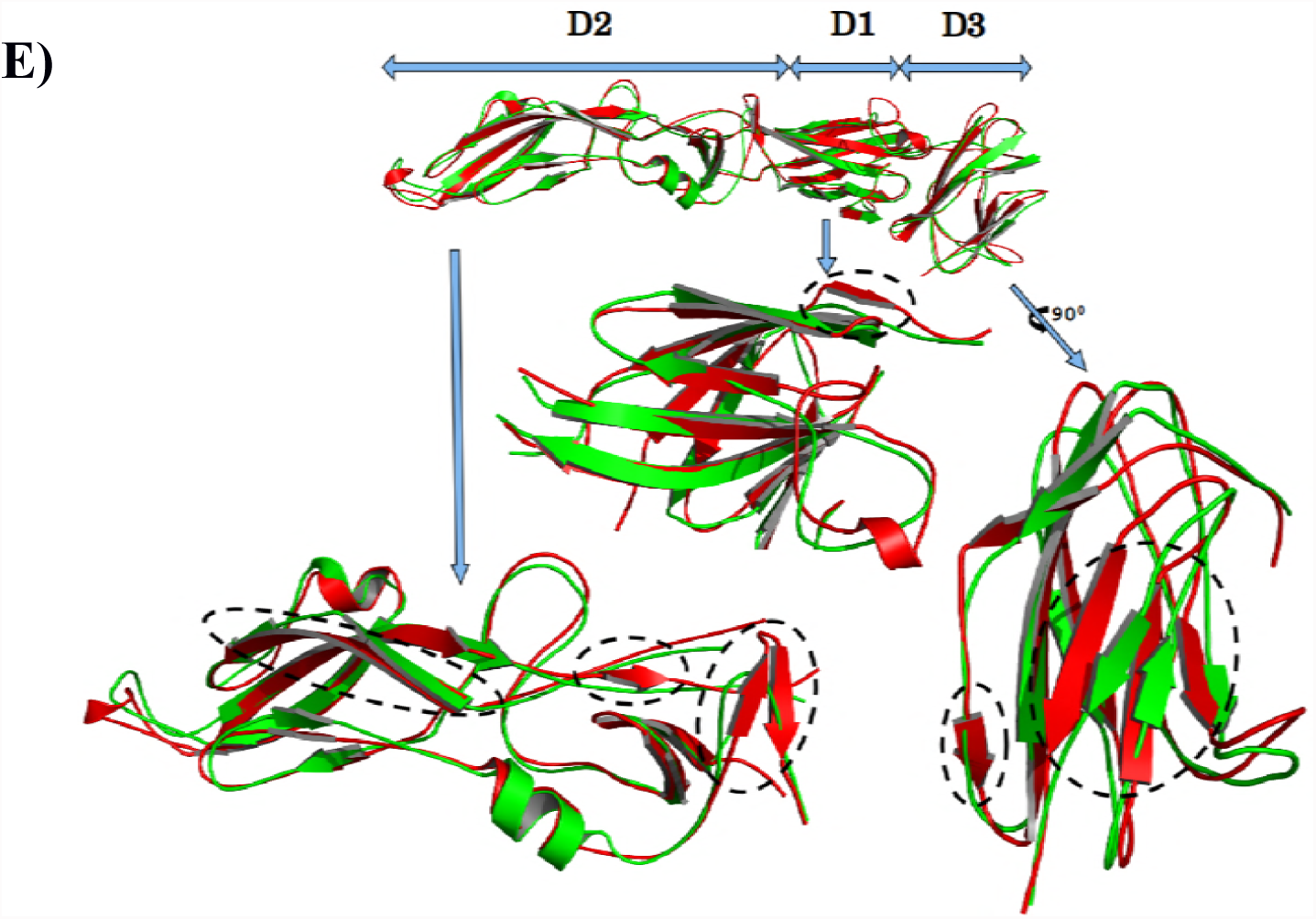
Residue-level energy contributions. Energy contributions at the three protein-protein interfaces - (A) A-A’, (B) B-A’, and (C) C-C in ZIKA (red) and DENV (black) from UA simulations, with domain-wise sequence alignments of two E proteins are depicted in insets. See Fig. 6D inset for E protein notations. **(D)** Protein structural network at a 5-fold vertex of ZIKA (left) and DENV (right). For clarity, five constituting E protein are colored differently. Color code for interactions - red: electrostatic, blue: H-bond, yellow: vdW. **(E)** Secondary structural content of the ZIKV (red) and DENV (green) E proteins. The regions displaying major differences are circled.

In consistence with the large inter-raft energy difference at B-A’ interface (see Fig. 6D), the number of ZIKA residue pairs contributing to this interface energy is found to be significantly larger than DENV as shown in Figure 7B. Among these, the residue pairs Lys340_r1_-Asp87_r2_, Thr353_r1_-Glu79_r2_, and Lys395_r1_-Asp83_r2_ in ZIKV showed large contribution, in contrary to the negligible interaction exhibited by the corresponding residue pairs Lys334_r1_-Asp87_r2_, His346_r1_-Glu79_r2_, and Glu386_r1_-Asn83_r2_ in DENV. While the sidechain-sidechain H-bond between Thr353_r1_ and Glu79_r2_ was very stable and persistent in ZIKV (see Fig. S6b for % H-bond persistence), the substituted His346_r1_ in DENV showed negligible interaction with Glu79_r2_. Similarly, the polar Lys395_r1_-Asp83_r2_ interaction in ZIKV was completely absent in DENV due to the sidechain-sidechain repulsion of substituted Glu386_r1_ with Asn83_r2_. Nonetheless, a few favorable interactions were present between the DENV residue pairs Ser300_r1_-Glu84_r2_, Lys334_r1_-Glu84_r2_, Met301_r1_-Glu84_r2_ etc. When compared with the EMdata, the H bonds formed by Ser298_r1_ and Glu338_r1_ respectively with Arg89_r2_ and Gln86_r2_ in DENV cryo-EM structure were retained only for 10% of simulation time (indicated by # in Fig. S6B, 7B). On the other hand, the cryo-EM structure H-bonds for ZIKA residue pairs Leu352 and Glu79, Thr353 and Glu79, Gly392 and Asp83 persisted >50% of the simulation time (Fig. S6B).

At C-C interface of ZIKA, the inter-raft stability was facilitated majorly by the residue pairs Glu393_r1_-Lys395_r2_ and Lys394_r1_-Glu393_r2_ (Fig. 7C). These residues involve in strong electrostatic interactions and contribute significantly toward the total inter-raft stabilization energy. In DENV, this sequence of charged residues EKK is substituted by PGQ, which fails to stabilize the 5-fold vertex efficiently, making it susceptible to temperature. Apart from these strong electrostatic interactions, the C-C interface in ZIKV is also stabilized by multiple H-bond interactions, in contrast to the weak van der Waals interactions in DENV. H-bond analysis at this interface shows persistent hydrogen bonds in ZIKV that match favorably well with the EM structure data. For example, H bonds found between Asn371_r1_ and Ser304_r2_, Thr313_r1_ and Ser304_r2_ in ZIKV cryo-EM structure were persistent for >60% of the simulation time (indicated by * in Fig. S6C). The cryo-EM H-bond between Val308_r1_ and Ser298_r2_ in DENV was also stable during the simulations. Interestingly, a recent structural study had proposed potential H-bond between Gln350_r1_ and Thr351_r2_ in CD-loops of the neighboring rafts as a potential contributor in ZIKA 5-fold vertex stability.(6) However, our data revealed <2% persistence of this H-bond during the entire simulation period. In fact, our result matches very well with a more recent mutagenesis study, which showed that alanine mutation at these residues Gln350 and Thr351 individually or in combination has no effect on ZIKA stability. Instead, our data suggests that the stability of ZIKV 5-fold vertex primarily stems from H-bonding interactions between the DE-loops (through residues Asn371 and Ser304) and electrostatic interactions between the FG-loops (due to EKK residue sequence).

A protein structural network often helps visualizing the complex interactions in protein-protein conjugates in a more tractable representation. Hence, we generated the structural network of a representative 5-fold vertex of ZIKA and DENV by combining the (significant) interactions at five adjoining C-C interfaces (i.e. five DIII domains at a 5-fold vertex). A protein structural network consists of nodes represented by aminoacids, connected to each other by edges that can be represented by different parameters. Studies have shown that the use of interaction energies as edge weights between nodes can efficiently capture the structural stability of protein-protein complexes.(20) Hence, we represent the ZIKA and DENV 5-fold vertices in the form of a network based on the pairwise interaction energies of their constituent aminoacids, and the results are shown in Fig. 7D. In the Figure, the Cα atoms of the interacting nodes are connected by edges representing their interactions, while the edge thicknesses represent the strength of interaction and color code represents the nature of interaction (red: electrostatic interactions >-5 kcal/mol, blue: H-bond interactions −2 to −5 kcal/mol, yellow: vdW interactions < −2 kcal/mol). It is evident form Fig. 7D that inter-raft communications in ZIKV are more robust through multiple electrostatic and H-bond interactions. Particularly, the intricate network of interlocking DE-, FG-loops among five DIIIs in ZIKA vertex was exceedingly robust that makes this envelope stable, even at high temperature or harsh conditions. On the other hand, the weaker communications in DENV, majorly assisted by feeble vdW interactions and few H-bonds, make this envelope prone to break.

Do the above-distinguished differences in two virus envelopes emerge, to some extent, from the intrinsic architecture of their constituent E proteins? In this context, we examined the secondary structural content of the envelope protein of both the viruses and the results are shown in Table S2 and Fig. 7E. As Table S2 clearly shows, ZIKA E protein contained > 10% secondary structure (ß-strand + Helix) than DENV. The E protein superimposed structures in Fig. 7E could easily distinguish these differences, domain-wise. It is evident form this Figure that all ZIKV domains, particularly domain II and III have significantly more antiparallel ß-sheets than DENV (highlighted by dotted circles). The scarcity of secondary structures in DENV was most prominent in the DIII domain, where majority of the ZIKA ß-sheets were replaced by coils in DENV. The implications of these changes lie on the fact that the longer beta strands in ZIKV could better stabilize the intermittent loops - CD-loop, DE-loop, FG-loop that involve in inter-raft interactions and, thus, making ZIKA 5-fold vertices very stable. On the other hand, more coil and less ß-sheet content in DENV introduce more dynamics/fluctuations in these structurally important loops, making DENV raft-raft interfaces weak and susceptible to temperature.

### Mutational studies revealed key residues responsible for greater ZIKA stability

From the above analyses, it became evident that strong association amongst key E protein residues at inter-raft interfaces was the primary reason for greater ZIKA stability. A more direct link of these residues on ZIKA envelope stability was further deciphered by performing computational mutagenesis studies, where the selected ZIKA residues were mutated to alanine. From the interaction energy data in Figure 7, we have identified the inter-raft residues Asp83, Asp87, Ser86, Glu133, Lys340, Gln344, Thr353, Glu393, Lys394, Lys395 for alanine mutations (Table S3). These identified residues were replaced with alanines and subsequently the whole virus envelope was subjected to thorough equilibration prior to 1µs long CG simulations, both at 29°C and 40°C. The visual inspection of the simulation trajectories revealed unstable envelope with destabilized vertices and inter-raft interfaces. We then performed RMSD and inter-raft angle analysis to capture the changes in detail. The RMSD plot in Fig. S7A clearly shows that a set of ZIKA rafts have undergone sharp increase in RMSD values within the first 100 ns of simulation run, indicating a loosened-up envelope even at 29°C. The inter-raft angle distributions also exhibited a disruption of the envelope by showing an asymmetric movements of the constituent rafts (Fig. S7B).

To locate the most crucial E protein residues responsible for greater ZIKA envelope stability, we subsequently examined a new system by mutating residues, Glu393, Lys394 and Lys395 only. These three specific residues were picked from the above list of ten mutations, noting their largest contribution in ZIKA inter-raft interactions (Fig. 7). The three residues were replaced by alanine and the whole virus envelope was simulated for 1µs using CG simulation protocol at 29°C. Interestingly, the ZIKV envelope displayed significantly weakened inter-raft interactions, as exemplified by the asymmetric angle distributions in Fig. S8. The 3D representation of the changes captures the breaking of 5-fold vertices with the formation of large holes, as shown in Fig. 8. This indicates that E protein residues Glu393, Lys394 and Lys395 at inter-raft interfaces play the most crucial role in ZIKV stability. As a validation of our mutagenesis results, we also simulated a control system by mutating residues Gln350 and Thr351 with alanines, following the experiment of Goo and co-workers.(21) This system was simulated both at 29°C and 40°C. Interestingly, our results show minimal effect of these mutations on ZIKA stability. The virus envelope remained intact at both temperatures, suggesting no significant H-bonding interaction between Gln350 and Thr351 in the parent system, as also was concluded by Goo and co-workers.

**Fig 8:**
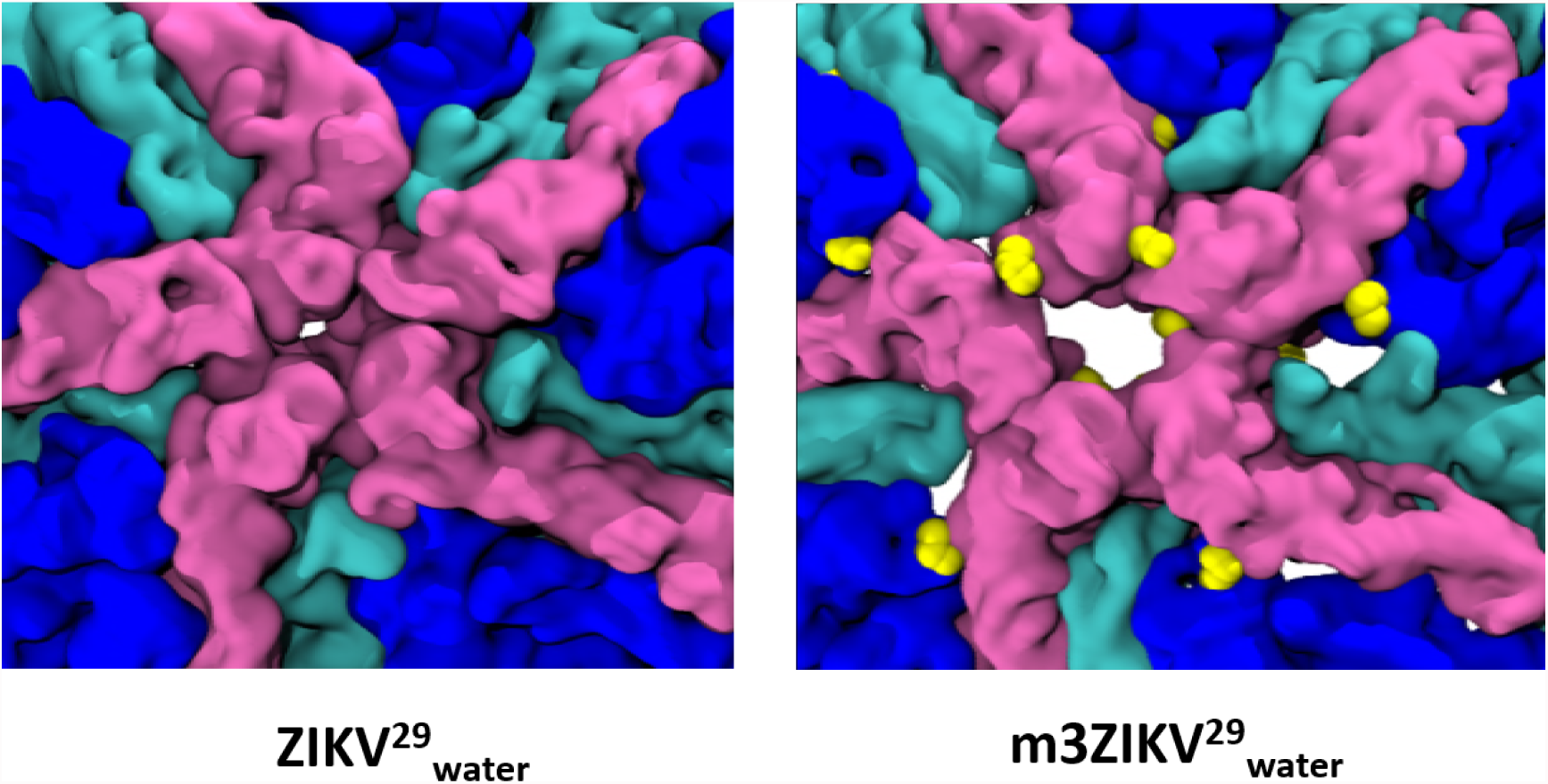
Specific mutations induced instability in ZIKV. Structure of 5-fold vertices of ZIKV^29^_water_ and mutated ZIKV^29^_water_ at the end of 1µs CG simulation. Mutated residues (E393A, K394A and K395A) are shown in yellow beads.

## CONCLUSIONS

ZIKA virus has spread all across the world very rapidly. Although the cryo-EM studies have revealed similar envelope structures of ZIKA and other flaviviruses, the more recent reports have suggested that only ZIKV can withstand high fever. In this study, we attempt to understand the underlying mechanism of differential stability of ZIKV and DENV2 at high temperatures. Our results from CG simulations show that, at these temperatures, ZIKV envelope retains its structural integrity, while DENV2 disintegrate through the raft-raft interfaces triggered by the formation of holes at the 5- and 3-fold vertices. In agreement with the cryo-EM maps at 37°C by Fibriansah et al., our MD simulations captured all class I – III DENV2 particles showing the loss of its icosahedral symmetry. (9)

To obtain finer details at the raft-raft interfaces, we then executed atomistic simulations. Results show that ZIKV vertices experience negligible changes with the constituting rafts maintaining strong contacts. Interactions at protein-level suggested that inter-raft interfaces are weaker than the intra-raft interfaces and intra-dimer association is the strongest among all pair interactions present on both ZIKA and DENV envelope surfaces, which is in good agreement with the experimental reports.(9) The protein structural network shows a stronger inter-raft communications in ZIKV through multiple electrostatic and H-bond interactions. Particularly, the intricate network of interlocking DE-loop and FG-loop among five DIII domains in ZIKA vertices was exceedingly robust that makes this envelope stable, even at high temperature. We found that the above-distinguished differences in two virus envelopes emerge, to some extent, from their intrinsic architecture of greater secondary structural content in ZIKV E protein than DENV. Our mutational results are validated by alanine mutations to the CD-loop residues Gln350 and Thr351 that showed no effect on ZIKA stability, in close accordance with a recent mutagenesis study.(21)

Recently, a set of monoclonal antibodies against various flaviviruses, including ZIKV has been tested.(22–25) These virus-neutralizing antibodies were primarily found to target the DIII domain of viral E protein. Interestingly, six of the ten ZIKV residues that we found crucial for ZIKV stability (Lys340, Gln344, Thr353, Glu393, Lys394, Lys395) belong to DIII domain. Thus, our study not only defines DE-, FG-, eE_0_-loops as potential ZIKV epitopes, but also provides detailed residue-label information that could pave way for designing specific ZIKV (and DENV2) antibodies. In addition, small molecule inhibitors binding to the identified E protein residues could be designed to inhibit E protein assembly/membrane fusion, similar to the recently reported HIV and DENV envelope-targeted inhibitors.(26–28)

## Methods

### System Preparation

Cryo-EM structures of ZIKV (PDBid: 5IRE) (5) and DENV envelopes (PDBid: 3J27)(7) at 3.8A and 3.6A resolution were obtained to prepare the initial conformations for simulations. The missing atoms and ZIKV E protein missing residues, 502:V-S-A:504 were incorporated using modeller 9v13 tool.(29) Similarly, the DENV M protein missing residues, 73: S-M-T:75 were included. The mutated ZIKV and DENV systems were also generated using modeller 9v13. The prepared systems were subjected to coarse-grained (CG) and united-atom (UA) simulations to explore their structural stability and inherent dynamics. The list of systems studied is shown in Table S1.

### Simulation details

Initially, we performed CG simulations of ZIKV and DENV envelopes to observe the large-scale conformational changes. We employed ElNeDyn force field by integrating the elastic network model with Martini CG parameters.(30) The elastic network model was employed to retain the secondary structures of viral E and M proteins during CG simulations. The intra and intermolecular interactions were described by standard Martini 2.2 force field (31) with four-to-one mapping, *i.e.* on average four heavy atoms and associated hydrogens were embedded to a single bead. In this model, four main types of beads - polar, non-polar, apolar, charged; and a number of subtypes were defined that could represent the chemical nature /atomistic description of the molecule in study. The systems were solvated in a cubic box by CG water, where four water molecules together is defined by a single bead. In the hollow systems, however, the water molecules inside the virus envelopes were removed. Periodic boundary conditions were applied in all directions. Default protonation states were assigned to all amino acids assuming neutral pH and counter ions were added to neutralize the systems. Systems were energy minimized using steepest descent algorithm. Simulations were performed at two different temperatures 29°C and 40°C. Pressure was maintained at 1 bar using Parrinello-Rahman barostat with isotropic pressure coupling. A cut-off of 1.2 nm was used for both electrostatic and van der Waals interactions. All the systems were extensively equilibrated upto 100ns by gradually reducing the positional restraints on backbone CG beads from 1000 kJ mol^-1^ nm^-2^ to 0 kJ mol^-1^ nm^-2^. The equilibrated structures were then subjected to production run of 4µs for ZIKV and 1µs for DENV with a time step of 20fs. All simulations were performed using Gromacs 5.0.7 simulation software.

Subsequently, two systems at 40°C were subjected to UA simulations using GROMOS96 53A6 forcefield.(32) Initially, the systems were briefly minimized using steepest descent and conjugate gradient algorithms, followed by solvation with explicit water (SPC model) in cubic periodic box. The salt concentration of 0.15M was maintained. The solvated systems were then subjected to extensive energy minimization, followed by thorough equilibration in NPT ensemble to adjust the solvent density. A cut off of 1.0 nm for both van der Waals and electrostatic interactions and particle mesh Ewald sum with real space cut-off at 1.0 nm were used. LINCS algorithm was used to constrain all bonds involving hydrogen atoms. The systems were equilibrated for 1 ns with a time step of 2 fs. Finally the production run was performed for 20 ns for each ZIKA and DENV system. We utilized gromacs trajectory analysis tools and in-house scripts to extract the informations from simulation data. Molecular mechanics generalized Born surface area (MMGBSA) method is used to calculate the inter- and intra-raft interaction energies, and subsequently residue-level decomposition was performed to obtain residue-residue interactions.(33)

### SUPPLEMENTAL INFORMATION

Supplemental Information includes eight figures, three tables, and two videos and can be found online.

### Author Contributions

Conceptualization, MHR, SS; Methodology, VRC, CP; Analysis, CP, VRC, MA, PDR, MHR; Investigation, VRC, CP, MA, SS; Writing-original Draft, CP; Writing-Review & Editing, SS; Supervision, SS.

## References

1. Zanluca C, Melo VCA de, Mosimann ALP, Santos GIV dos, Santos CND dos, Luz K. 2015. First report of autochthonous transmission of Zika virus in Brazil. Mem Inst Oswaldo Cruz 110:569–572.

2. Martines RB, Bhatnagar J, Keating MK, Silva-Flannery L, Muehlenbachs A, Gary J, Goldsmith C, Hale G, Ritter J, Rollin D, Shieh W-J, Luz KG, Ramos AM de O, Davi HPF, Kleber de Oliveria W, Lanciotti R, Lambert A, Zaki S. 2016. *Notes from the Field*: Evidence of Zika Virus Infection in Brain and Placental Tissues from Two Congenitally Infected Newborns and Two Fetal Losses — Brazil, 2015. MMWR Morb Mortal Wkly Rep 65:1–2.

3. Foy BD, Kobylinski KC, Chilson Foy JL, Blitvich BJ, Travassos da Rosa A, Haddow AD, Lanciotti RS, Tesh RB. 2011. Probable non-vector-borne transmission of Zika virus, Colorado, USA. Emerg Infect Dis 17:880–2.

4. Lindenbach BD, Rice CM. 2007. Flaviviridae: The Viruses and Their Replication. Fields Virol 1101–1151.

5. Sirohi D, Chen Z, Sun L, Klose T, Pierson TC, Rossmann MG, Kuhn RJ. 2016. The 3.8 Å resolution cryo-EM structure of Zika virus. Science 352:467–70.

6. Kostyuchenko VA, Lim EXY, Zhang S, Fibriansah G, Ng T-S, Ooi JSG, Shi J, Lok S-M. 2016. Structure of the thermally stable Zika virus. Nature 533:425–428.

7. Zhang X, Ge P, Yu X, Brannan JM, Bi G, Zhang Q, Schein S, Zhou ZH. 2013. Cryo-EM structure of the mature dengue virus at 3.5-Å resolution. Nat Struct Mol Biol 20:105–110.

8. Mansuy JM, Dutertre M, Mengelle C, Fourcade C, Marchou B, Delobel P, Izopet J, Martin-Blondel G. 2016. Zika virus: high infectious viral load in semen, a new sexually transmitted pathogen? Lancet Infect Dis 16:405.

9. Fibriansah G, Ng T-S, Kostyuchenko VA, Lee J, Lee S, Wang J, Lok S-M. 2013. Structural Changes in Dengue Virus When Exposed to a Temperature of 37 C. J Virol 87:7585–7592.

10. Lim X-X, Chandramohan A, Lim XYE, Bag N, Sharma KK, Wirawan M, Wohland T, Lok S-M, Anand GS. 2017. Conformational changes in intact dengue virus reveal serotype-specific expansion. Nat Commun 8:14339.

11. Freddolino PL, Arkhipov AS, Larson SB, McPherson A, Schulten K. 2006. Molecular Dynamics Simulations of the Complete Satellite Tobacco Mosaic Virus. Structure 14:437–449.

12. Arkhipov A, Freddolino PL, Schulten K. 2006. Stability and Dynamics of Virus Capsids Described by Coarse-Grained Modeling. Structure 14:1767–1777.

13. Perilla JR, Schulten K. 2017. Physical properties of the HIV-1 capsid from all-atom molecular dynamics simulations. Nat Commun 8:15959.

14. Reddy T, Shorthouse D, Parton DL, Jefferys E, Fowler PW, Chavent M, Baaden M, Sansom MSP. 2015. Nothing to Sneeze At: A Dynamic and Integrative Computational Model of an Influenza A Virion. Structure 23:584–597.

15. Larsson DSD, Liljas L, van der Spoel D. 2012. Virus Capsid Dissolution Studied by Microsecond Molecular Dynamics Simulations. PLoS Comput Biol 8:e1002502.

16. Reddy T, Sansom MSP. 2016. The Role of the Membrane in the Structure and Biophysical Robustness of the Dengue Virion Envelope. Structure 24:375–382.

17. Marzinek JK, Holdbrook DA, Huber RG, Verma C, Bond PJ. 2016. Pushing the Envelope: Dengue Viral Membrane Coaxed into Shape by Molecular Simulations. Structure 24:1410–1420.

18. Katen S, Zlotnick A. 2009. Chapter 14 The Thermodynamics of Virus Capsid Assembly, p. 395–417. *In* Methods in enzymology.

19. Ceres P, Zlotnick A. 2002. Weak protein-protein interactions are sufficient to drive assembly of hepatitis B virus capsids. Biochemistry 41:11525–31.

20. Appadurai R, Senapati S. 2016. Dynamical Network of HIV-1 Protease Mutants Reveals the Mechanism of Drug Resistance and Unhindered Activity. Biochemistry 55:1529–1540.

21. Goo L, Dowd KA, Smith ARY, Pelc RS, DeMaso CR, Pierson TC. 2016. Zika Virus Is Not Uniquely Stable at Physiological Temperatures Compared to Other Flaviviruses. MBio 7:e01396–16.

22. Crill WD, Roehrig JT. 2001. Monoclonal antibodies that bind to domain III of dengue virus E glycoprotein are the most efficient blockers of virus adsorption to Vero cells. J Virol 75:7769–73.

23. Beasley DWC, Barrett ADT. 2002. Identification of neutralizing epitopes within structural domain III of the West Nile virus envelope protein. J Virol 76:13097–100.

24. Zhao H, Fernandez E, Dowd KA, Speer SD, Platt DJ, Gorman MJ, Govero J, Nelson CA, Pierson TC, Diamond MS, Fremont DH. 2016. Structural Basis of Zika Virus-Specific Antibody Protection. Cell 166:1016–1027.

25. Yang M, Dent M, Lai H, Sun H, Chen Q. 2017. Immunization of Zika virus envelope protein domain III induces specific and neutralizing immune responses against Zika virus. Vaccine 35:4287–4294.

26. Lin P-F, Blair W, Wang T, Spicer T, Guo Q, Zhou N, Gong Y-F, Wang H-GH, Rose R, Yamanaka G, Robinson B, Li C-B, Fridell R, Deminie C, Demers G, Yang Z, Zadjura L, Meanwell N, Colonno R. 2003. A small molecule HIV-1 inhibitor that targets the HIV-1 envelope and inhibits CD4 receptor binding. Proc Natl Acad Sci 100:11013–11018.

27. Schmidt AG, Lee K, Yang PL, Harrison SC. 2012. Small-Molecule Inhibitors of Dengue-Virus Entry. PLoS Pathog 8:e1002627.

28. Modis Y, Ogata S, Clements D, Harrison SC. 2003. A ligand-binding pocket in the dengue virus envelope glycoprotein. Proc Natl Acad Sci U S A 100:6986–91.

29. Webb B, Sali A. 2016. Comparative Protein Structure Modeling Using MODELLER, p. 5.6.1–5.6.37. *In* Current Protocols in Bioinformatics. John Wiley & Sons, Inc., Hoboken, NJ, USA.

30. Periole X, Cavalli M, Marrink S-J, Ceruso MA. 2009. Combining an Elastic Network With a Coarse-Grained Molecular Force Field: Structure, Dynamics, and Intermolecular Recognition. J Chem Theory Comput 5:2531–2543.

31. Marrink S-J, Risselada H-J, Yefimov S, Tieleman D-P, Vries A-H de. 2007. The MARTINI Force Field: Coarse Grained Model for Biomolecular Simulations. J Phys Chem B 111:7812–7824.

32. Oostenbrink C, Villa A, Mark AE, Van Gunsteren WF. 2004. A biomolecular force field based on the free enthalpy of hydration and solvation: The GROMOS force-field parameter sets 53A5 and 53A6. J Comput Chem 25:1656–1676.

33. Mongan J, Simmerling C, McCammon J-A, Case D-A, Onufriev A. 2006. Generalized Born Model with a Simple, Robust Molecular Volume Correction. J Chem Theory Comput 3:156–169.

